# Cardiomyopathy-Associated Variants Alter the Structure and Function of the α-Actinin-2 Actin-Binding Domain

**DOI:** 10.1101/2023.05.09.539883

**Authors:** Alexandra E Atang, Robyn T. Rebbeck, David D. Thomas, Adam W. Avery

**Affiliations:** Department of Chemistry, Oakland University, Rochester, MI 48309-4479, USA; Department of Biochemistry, Molecular Biology and Biophysics, University of Minnesota, Minneapolis 55455, USA

**Keywords:** Actin binding, circular dichroism, thermal denaturation, protein folding

## Abstract

Hypertrophic cardiomyopathy (HCM), dilated cardiomyopathy (DCM), and restrictive cardiomyopathy (RCM) are characterized by thickening, thinning, or stiffening, respectively, of the ventricular myocardium, resulting in diastolic or systolic dysfunction that can lead to heart failure and sudden cardiac death. Recently, variants in the *ACTN2* gene, encoding the protein α-actinin-2, have been reported in HCM, DCM, and RCM patients. However, functional data supporting the pathogenicity of these variants is limited, and potential mechanisms by which these variants cause disease are largely unexplored. Currently, NIH ClinVar lists 34 *ACTN2* missense variants, identified in cardiomyopathy patients, which we predict are likely to disrupt actin binding, based on their localization to specific substructures in the α-actinin-2 actin binding domain (ABD). We investigated the molecular consequences of three ABD localized, HCM-associated variants: A119T, M228T and T247M. Using circular dichroism, we demonstrate that the mutant ABD proteins can attain a well-folded state. However, thermal denaturation studies show that all three mutations are destabilizing, suggesting a structural disruption. Importantly, A119T decreased actin binding, and M228T and T247M cause increased actin binding. We suggest that altered actin binding underlies pathogenesis for cardiomyopathy mutations localizing to the ABD of α-actinin-2.

## 1. Introduction

In the United States, the prevalence of heart failure due to cardiomyopathy was estimated at 6.7 million cases in 2020, an increase of 12% over 2 years.[1] Two of the most common cardiomyopathies are Dilated Cardiomyopathy (DCM) and Hypertrophic Cardiomyopathy (HCM), combined estimated to occur in greater than 0.2% of the general population[2, 3]. Restrictive Cardiomyopathy (RCM), a less common form, is estimated to account for 5% of total cardiomyopathy cases[4]. These disorders are characterized by thickening (HCM), thinning (DCM), or stiffening (RCM) of the ventricular myocardium[5]. These alterations to the ventricle wall cause diastolic or systolic dysfunction, and can lead to heart failure and sudden cardiac death[6]. A better understanding of cardiomyopathy etiology and pathogenesis is thus important for reducing the burden cardiac disease for a significant portion of the population.

The familial forms of DCM, HCM, and RCM typically display autosomal dominant inheritance[5]. Interestingly, dominant mutations in the *ACTN2* gene encoding the Z-disk protein α-actinin-2 have been reported in DCM[7], HCM[8] and RCM[9] patients. How different variants in the *ACTN2* gene give rise to different cardiomyopathies is unclear. Currently, the NIH ClinVar database[10] lists 324 cardiomyopathy-associated *ACTN2* variants that result in amino acid changes in the encoded α-actinin-2 protein. The pathogenicity of two *ACTN2* missense variants, A119T and M228T, is supported by co-segregation of the variants with heart disease across multiple generations within families[8, 11]. However, for most reported *ACTN2* variants, the pedigree is either too small, or no family history is available to conclude that a genetic variant is disease causing. In these cases, additional functional studies are needed to determine if a variant is detrimental to α-actinin-2 function.

Alpha-actinin-2 is expressed in skeletal and cardiac muscle fibers[12], and is required for normal sarcomere organization in the heart[13]. Within the Z-disk, α-actinin-2 functions to cross-link antiparallel actin filaments from neighboring sarcomeres. Alpha-actinin-2 contains an N-terminal actin-binding domain (ABD; amino acids 1-259) consisting of two calponin homology subdomains (CH1 and CH2). Assembly of the Z-disk requires α-actinin-2 binding to actin. Interestingly, α-actinin-2 localization to the Z-disk is highly dynamic[14-16]. This suggests that dynamic binding of α-actinin-2 to F-actin is required in healthy tissue to enable rapid Z-disk remodeling.

Alpha-actinin is the ancestral member of the spectrin superfamily of cytoskeletal proteins that includes the human disease-associated proteins: β-III-spectrin; α-actinin-1 and -4; filamin A, B, and C; and dystrophin[17-26]. We previously characterized a β-III-spectrin L253P missense mutation that is localized to the ABD and causes the neurological disorder spinocerebellar ataxia type 5 (SCA5). In vitro co-sedimentation assays revealed that L253P causes a striking 1000-fold increase in actin affinity[18]. Further, our cryo-EM structure of F-actin complexed with the L253P ABD demonstrated that actin binding requires a conformational “opening” at the interface of the two CH domains that comprise the ABD. This enables CH1 to bind actin aided by the previously uncharacterized, unstructured N-terminal region that becomes α-helical upon binding[27]. Strikingly, this N-terminal helix preceding the conserved CH1 domain is required for association with actin, as truncation eliminates binding. Alpha-actinin-2 likely binds actin by a similar mechanism. Indeed, our review of cryo-EM data for the α-actinin-4-actin complex[28] suggested that the N-terminus of α-actinin-2 also binds actin[27].

However, mechanistic studies of cardiomyopathy-associated variants in the α-actinin-2 ABD are scarce. Haywood et al, reported that the HCM-associated variant A119T caused protein destabilization *in vitro*, aggregation of the mutant α-actinin-2 in cardiomyocytes, and reduced Z-disk incorporation[15]. A119T was reported to cause a 2-fold decrease in actin-binding affinity[15], although the reported decrease in affinity was not statistically significant. A study of an HCM-associated variant, T247M, established in iPSC-derived cardiomyocytes, showed that the variant is disruptive at the cellular level, inducing HCM phenotypes, including Z-disk myofibrillar disarray and disrupted contractility[16]. Moreover, T247M, like A119T, caused α-actinin-2 aggregation and reduced incorporation into Z-disks [29]. How the T247M variant impacts actin binding is not known. A recent study has shown that homozygous expression of HCM associated M228T variant in mice is embryonic lethal[30]. Further, low α-actinin-2 mutant protein levels suggested that the mutant promotes destabilization contributing to increased proteolysis[30]. Because of the dynamic nature of α-actinin-2 at the Z-disk, and the requirement of actin binding for α-actinin-2 incorporation into the Z-disk, it is likely that ABD variants that disrupt actin binding will adversely impact cardiomyocyte function.

In this study, we report our analyses of the structural position of numerous cardiomyopathy-associated *ACTN2* missense variants to assess the potential of the variants to disrupt α-actinin-2 actin binding. We further report our characterization of the structural and functional consequences of select missense variants localized to different substructures in the alpha-actinin-2 ABD.

## 2. Materials and Methods

### 2.1 Structural analysis

Analyses were performed using a crystal structure of the wild-type alpha-actinin-2 ABD (PDB:5A36)[15]. Actin binding region identified by homology comparison with three different ABDs including of β-III-spectrin, filamin A and utrophin ABD-actin complexes[27, 31, 32]. Analyses of the structure, including identification of predicted contacts for different mutated residues, was performed with PyMOL v2.5.4

### 2.2 Protein expression and purification

The coding sequence for α-actinin-2-ABD was obtained from Addgene (RefSeq: NM_001103.3, Plasmid ID 52669) and PCR amplified using the forward primer AAACACCTGCAAAAAGGTATGAACCAGATAGAGCCCGGC and reverse primer AAATCTAGATTACTCCGCGCCCGCAAAAGCGTG. The PCR product was digested with restriction enzymes AarI and XbaI and ligated into BsaI digested, pE-SUMOpro (LifeSensors). Mutations were introduced into the generated pE-SUMO-α-actinin-2 ABD through site-directed PCR mutagenesis (PfuUltra High-Fidelity DNA Polymerase, Agilent). The following primers were used to introduce cardiomyopathy mutations into the α-actinin-2 ABD sequence.

A119T forward: GTGAAACTGGTGTCCATCGGCACTGAAGAAATTGTTGATGGCAAT

A119T reverse: ATTGCCATCAACAATTTCTTCAGTGCCGATGGACACCAGTTTCAC

M228T forward: AGCACCTGGATATTCCTAAAACGTTGGATGCTGAAGACATCGTG

M228T reverse: CACGATGTCTTCAGCATCCAACGTTTTAGGAATATCCAGGTGCT

T247M forward: CCCGATGAAAGAGCCATCATGATGTACGTCTCTTGCTTCTACCAC

T247M reverse: GTGGTAGAAGCAAGAGACGTACATCATGATGGCTCTTTCATCGGG

Sequence verified mutant and wild-type DNAs were transformed into Rosetta 2 (DE3) *E. coli* (NOVAgen.1 L bacteria cultures in LB broth containing ampicillin (100 μg/mL) and chloramphenicol (34 μg/mL) were grown at 37 °C. Upon OD_550_ of 0.50, ABD protein expression was induced with 0.5 mM IPTG for 6 hours at 300 RPM shaking at room temperature. Bacteria cultures were pelleted at 2987 RCF for 30 min at 4 °C, and pellets stored at -20°C until further use. Bacterial pellets were prepared for lysis through resuspension in lysis buffer (50 mM Tris pH 7.5, 300 mM NaCl, 25% sucrose, and protease inhibitors (Complete Protease Inhibitor tablet, EDTA-free, Roche)), and incubated with lysozyme (Sigma) for 1 h at 4 °C, followed by a freeze-thaw cycle in an isopropanol-dry ice bath. To the lysate, MgCl_2_ (10 mM final concentration) and DNase1 (Roche) (8 U/mL final concentration) were added and stirred slowly for 1 h at 4 °C. Lysates were clarified at 18,000 RPM at 4 °C for 30 min, in a Sorvall SS-34 rotor. Supernatants were decanted and syringe filtered through 0.45 μm disk filters. Poly-Prep (Biorad) chromatography columns containing Ni-NTA agarose (Qiagen) were equilibrated in binding buffer (50 mM Tris pH 7.5, 300 mM NaCl and 20 mM imidazole) and filtrate was loaded. Binding buffer was used to wash the columns and ABD proteins were eluted with elution buffer((50 mM Tris pH 7.5, 300 mM NaCl and 150 mM imidazole). Elution fractions for each ABD protein were pooled and loaded into a Slide-a-Lyzer, 10 K MWCO, dialysis cassette (ThermoScientific) and dialysis carried out overnight in SUMO buffer (25 mM Tris pH 7.5, 150 mM NaCl, 5 mM β-mercaptoethanol), at 4 °C. Cleavage of the 6X-His-SUMO tag from the ABD protein was performed using 1:10 mass ratio of Ulp1 SUMO protease:ABD in a 1 h incubation at 4 °C. Poly-Prep chromatography columns containing 1 mL Ni-NTA agarose equilibrated in SUMO buffer were used to remove cleaved 6X-His SUMO tags and His-tagged SUMO protease from the ABD proteins. Eluted fractions containing ABD proteins were pooled for each ABD and loaded into a Slide-a-Lyzer, 10 K MWCO, dialysis cassette for dialysis in storage buffer (10 mM Tris pH 7.5, 150 mM NaCl, 2 mM MgCl2, and 1 mM DTT), at 4 °C overnight. Purified ABD proteins were recovered and measured for concentration using Bradford assay (Biorad).ABD proteins were snap frozen in liquid nitrogen and stored at -80 °C for future use.

### 2.3 Circular dichroism measurements

Previously purified ABD proteins were thawed and clarified at 43,000 RPM at 4 °C for 30 min, in a Beckman TLA 100.3 rotor. Bradford assay was used to determine the ABD protein concentrations. ABD proteins were diluted to 150 – 200 ng/μL in storage buffer. CD spectra and thermal denaturation studies were performed with a Jasco J-815 Spectropolarimeter with a Peltier temperature controller. Baseline correction scans were obtained using the storage buffer immediately prior to analysis. Analyses ranged between 200 and 260 nm at 25 °C. Immediately following wavelength scans, unfolding was measured at 222 nm with steadily ramping temperatures from 20 – 85 °C. Using Prism 9 (GraphPad), a nonlinear regression analysis was performed to determine melting temperature for two-state and three state transitions[33]. The two-state unfolding is fit by equation 1,

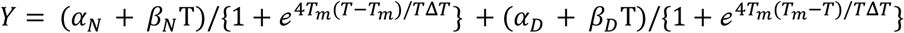

where α_N_, β_N_, α_D_, and β_D_ are the intercepts and slopes of the respective native and denatured states, Y is the signal at 222 nm at temperature T, ΔT is the unfolding transition width and T_m_ is the melting temperature. The three-state unfolding is fit by the expanded equation 2,

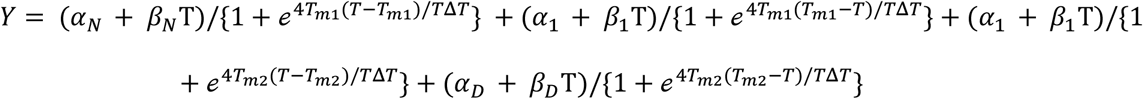

where, α_1_ and β_1_ are the intercept and slope of the intermediate state, T_m1_ and T_m2_ are the respective melting temperatures of the first state between folded and the intermediate and the second state between the intermediate and unfolded state. The four-state unfolding is fit by the expanded equation 3,

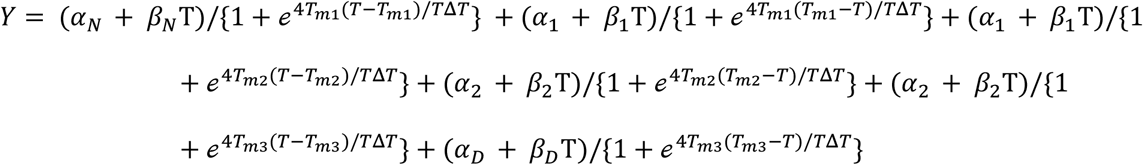

where, α_2_ and β_2_ are intercept and slope of the second intermediate state, T_m2_ and T_m3_ are the melting temperatures of the second state between the first and second intermediate and the second intermediate and the unfolded state.

### 2.4 F-actin co-sedimentation assays

Purified F-actin was prepared from rabbit skeletal muscle (Pel-Freez Biologics) as previously described[34]. Purified ABD proteins were thawed and clarified as described above. F-actin and ABD protein concentrations were determined though Bradford assay. As performed previously[34], binding reactions were prepared with 2 μM ABD protein and F-actin concentrations ranging from 3 – 120 μM in F-buffer (10 mM Tris pH 7.5, 150 mM NaCl, 0.5 mM ATP, 2 mM MgCl2, and 1 mM DTT). Immediately following 30 min incubation in ambient room temperature, binding reactions were centrifuged at 50,000 RPM for 30 min at 25 °C in a TLA-100 rotor (Beckman). Following centrifugation, supernatants were immediately sampled and mixed with 4X Laemmle sample buffer (Biorad). Samples of supernatant were separated by SDS-PAGE and resulting bands were visualized using Coomassie Brilliant Blue R-250 (Biorad) solution, followed by background removal with destain solution. An Azure Sapphire imager on the 680 nm channel was used to document the gels. The generated image files allowed quantification of ABD protein band fluorescence intensities using Image Studio Lite version 5.2 software. A standard curve was generated using incremental amounts of ABD proteins (0-2 μM) on a similarly treated gel, relating supernatant ABD fluorescence intensity to known ABD concentrations. As previously described [18], dissociation constants (Kd) were calculated using Prism 9 (Graphpad) through a nonlinear regression fit utilizing a one-site specific binding equation, constraining Bmax to 1.

## 3. Results

### 3.1 Cardiomyopathy missense variants localize to actin-binding surfaces or the CH1-CH2 interface in the α-actinin-2 ABD

We analyzed 71 cardiomyopathy-associated *ACTN2* variants in NIH ClinVar to evaluate the potential of the variants to disrupt actin binding, based on their localization to specific substructures in the ABD. We mapped predicted actin-binding surfaces in the crystal structure of the wild-type α-actinin-2 ABD (PDB: 5A36)[15] based on homology to the actin-binding surfaces identified in cryo-EM models of β-III-spectrin, filamin A and utrophin ABD-actin complexes[27, 31, 32], **Figure 1**. In addition, residues mediating interaction of CH1 to the regulatory CH2 subdomain were identified using Pymol. Of the 71 ClinVar variants in the ABD, 34 localize to actin-binding surfaces in CH1 or the N-terminus, or to the CH1-CH2 interface. We predict that variants localizing to actin-binding surfaces will decrease actin binding. In contrast, variants localizing to the CH1-CH2 interface are likely to increase actin binding by alleviating CH2 occlusion of CH1 binding to actin.

**Figure 1.**
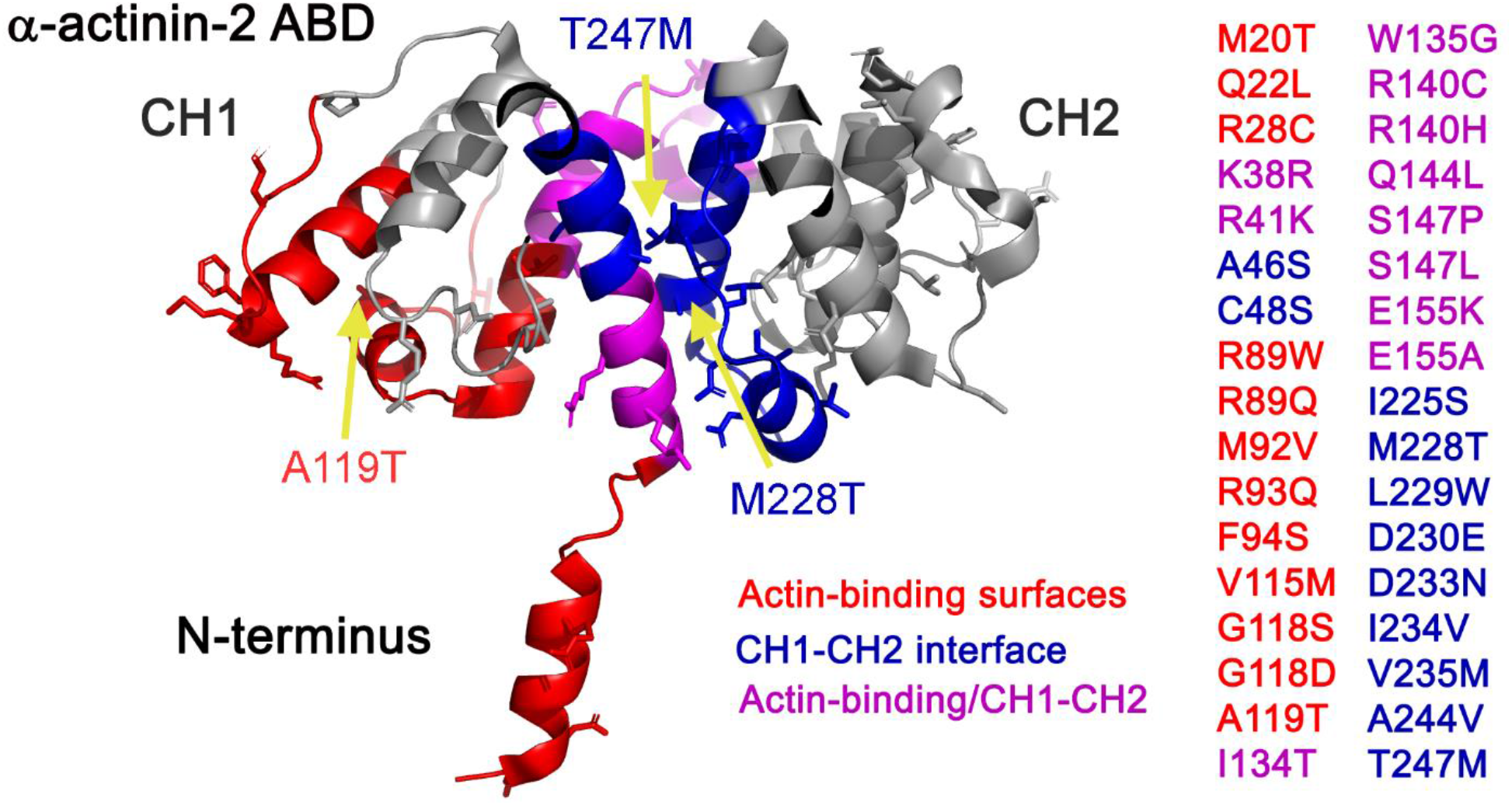
Mapping of *ACTN2* variants in the ABD. The α-actinin-2 ABD crystal structure (PDB: 5A36)[15] with actin-binding surfaces (red), contact regions between CH1 and CH2 subdomains (blue), and regions involved in both interactions (magenta). Listed are 34 cardiomyopathy missense variants localizing to these surfaces (sidechains of native residues shown). Yellow arrows indicate positions of A119T, M228T and T247M.

### 3.2 Human α-actinin-2 cardiomyopathy missense variants attain folded states but lose cooperative protein unfolding

To assess the impact of the variants on ABD structure, circular dichroism (CD) spectroscopy was performed. The CD absorption spectra for the wild-type ABD shows minima at 208 and 222 nm, an absorption profile characteristic of an alpha-helical protein, **Figure 2**. Similarly, the three variants showed nearly identical absorption profiles. This indicates the variant ABDs are well-folded like wild-type. CD was further used to monitor the unfolding of the ABD proteins as a function of temperature. The wild-type protein displayed cooperative unfolding, with a single step, two-state transition, and a melting temperature (T_m_) of 63.5°C, **Figure 3**. This T_m_ is similar a previously reported T_m_ (62.3°C) for a wild-type α-actinin-2 ABD construct containing amino acids 19-266[15]. In contrast, the melt curves for all mutants showed unfolding at lower temperatures and a clear loss in cooperative unfolding. The melt curve for A119T showed two transitions that were fit with two T_m_ values: T_m1_ = 60.9°C and 71.9°C (average T_m_’s from duplicate melt curves). Duplicate melting curves were nearly identical, supporting the accuracy of the melt profile. Interestingly, M228T and T247M mutants unfolded with very similar melt curves. These melt curves showed three transitions. For M228T, the three T_m_ values were 51.3°C, 60.7°C and 72.3°C. For T247M, the T_m_ values were similar: 50.6°C, 61.5°C and 72.0°C. The loss in cooperativity for M228T and T247M suggests reduced physical coupling between CH subdomains, and is consistent with our model that increased actin binding is associated with opening of the CH1/CH2 interface.

**Figure 2.**
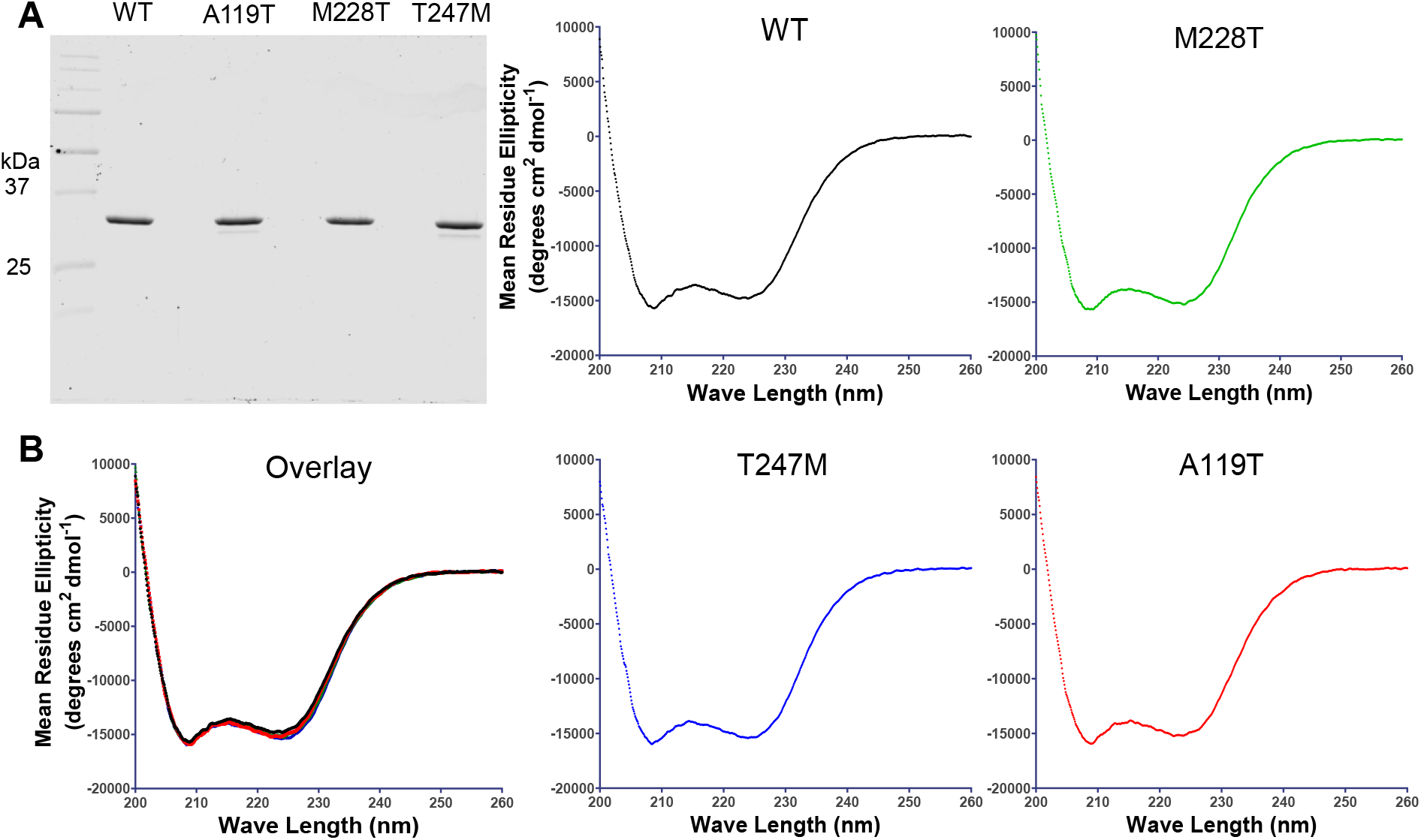
α-actinin-2 mutants attain secondary structures similar to wild-type. A. Coomassie blue stained gel image showing purified wild-type (WT) and mutant ABD proteins. All ABD proteins ran at the predicted size of 30 kDa. B. Circular dichroism absorption spectra show α-helical profiles for wild-type and mutant ABDs. CD spectra between 200 nm and 260 nm for individual WT, A119T, M228T, and T247M proteins. Minima at 208 and 222 nm are characteristic of α-helical fold.

**Figure 3.**
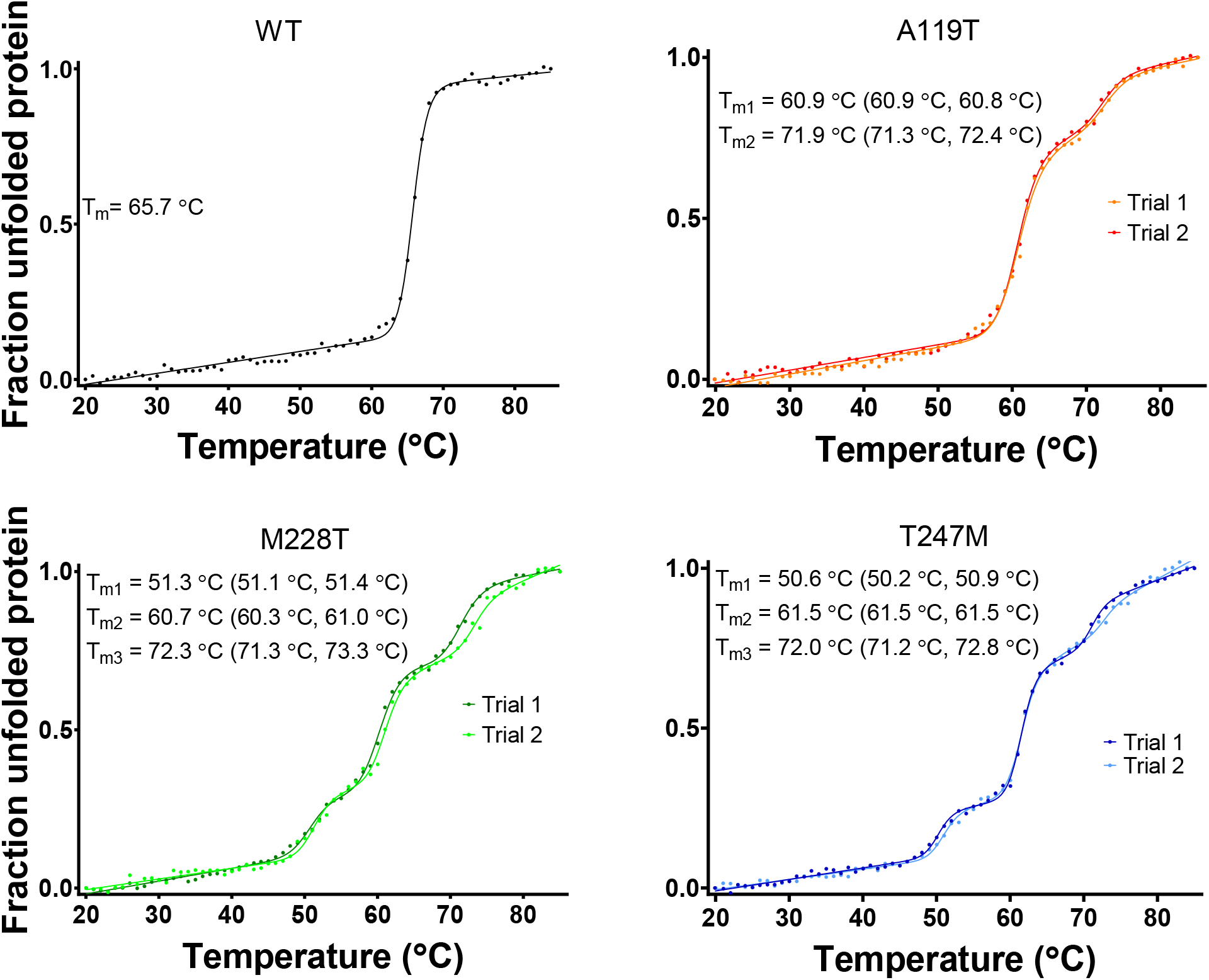
Thermal denaturation curves showing that mutant ABD proteins exhibit a loss of cooperative unfolding. Melting curves of purified ABD proteins, obtained by observing circular dichroism absorption at 222 nm while heating sample from 20 °C to 85 °C. The cooperative, two-state, transition is observed for wild-type ABD. A three-state transition is observed for A119T ABD in trial 1 (orange) and trial 2 (red). A four-state transition is observed for M228T ABD; trial 1 (dark green) and trial 2 (light green) and T247M ABD; trial 1 (dark blue) and trial 2 (light blue). For each transition, the average T_m_ from duplicate melts is indicated, followed by the individual replicate T_m_ values in parentheses.

### 3.3 Human α-actinin-2 cardiomyopathy missense variants disrupt actin binding

To test our prediction that CH1-CH2 interface variants in α-actinin-2 will increase actin binding, actin co-sedimentation assays were performed for the variants M228T and T247M. We additionally tested the A119T variant that localizes to an actin binding surface, and is thus predicted to decrease actin binding. The wild-type ABD showed dose responsive binding with increasing F-actin concentration, **Figure 4**. The average dissociation constant (Kd) of the wild-type ABD was 98 ± 7 μM. Significantly, both M228T and T247M ABDs were entirely bound to actin over the entire range of actin concentrations. This indicates that both M228T and T247M mutant ABDs bind actin with much higher affinity than wild-type, with Kd values that must be submicromolar. In contrast, A119T resulted in reduced binding of the ABD to actin. The Kd value of A119T was 466 ± 90 μM, a statistically significant reduction in the equilibrium binding constant relative to wild-type. Thus, our predictions for the impact of A119T, M228T and T247M on the binding of the ABD to actin were correct.

**Figure 4.**
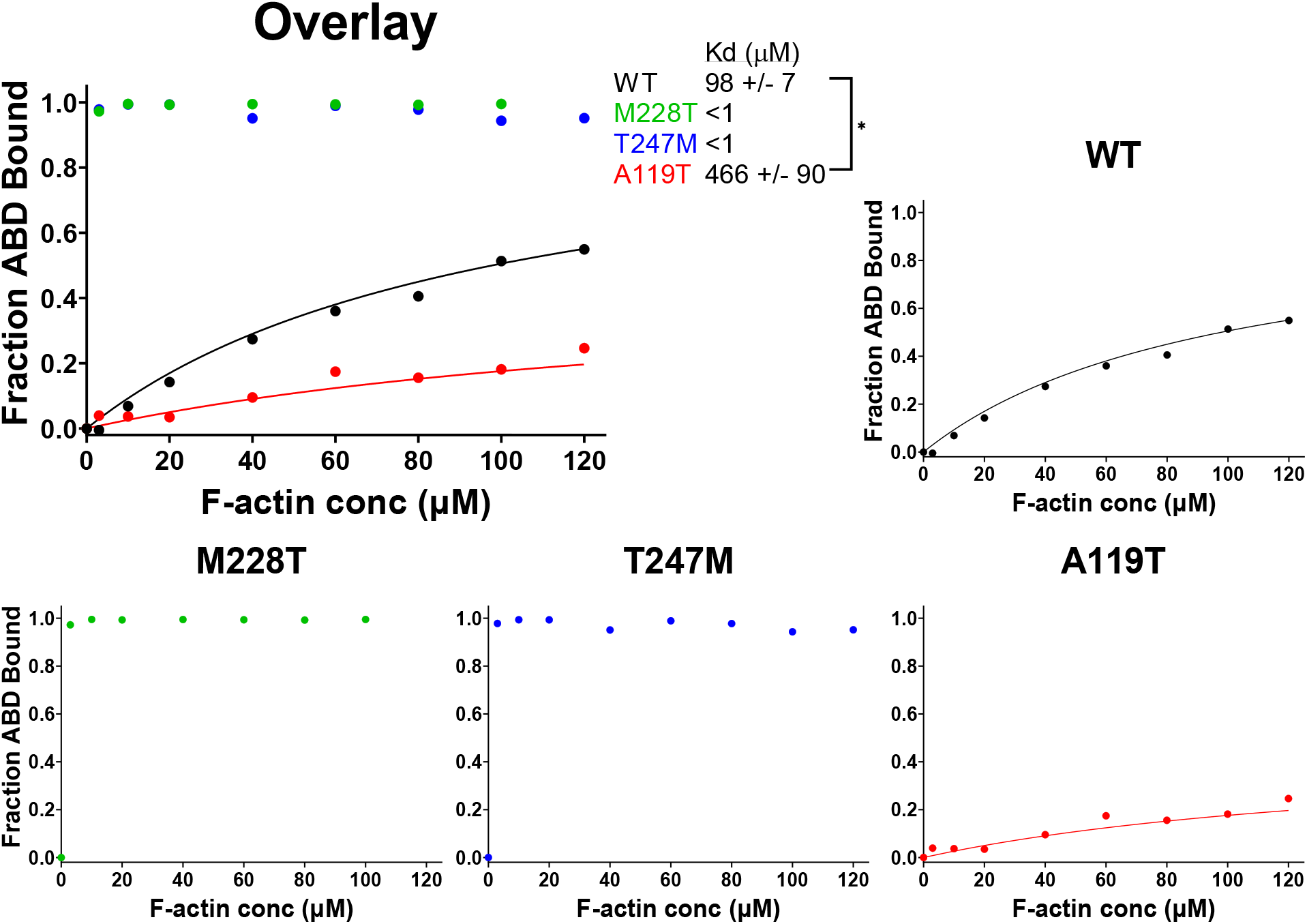
Actin co-sedimentation assays showing that ABD mutations cause aberrant actin binding. Cosedimentation of M228T and T247M ABD proteins show increased actin binding compared to wild-type. Cosedimentation of A119T ABD protein shows decreased actin binding compared to wild-type. Representative binding data and curve fit is shown for wild-type and A119T ABD proteins. At 2 μM, M228T and T247M ABD proteins are completely bound to actin. Representative binding data is shown for M228T and T247M. Kd value shown as mean ±SD, n = 4-6 binding assays.

## 4. Discussion

Many cardiomyopathy-associated missense variants have been reported in the *ACTN2* gene. However, the molecular consequences of these variants in the encoded α-actinin-2 protein are largely unexplored. Here we focused on variants that localize to the actin-binding domain (ABD) of α-actinin-2. Of 71 ABD-localized variants, 34 are positioned at actin-binding surfaces in CH1 or the N-terminus, or the CH1/CH2 interface. Importantly, we showed that two CH1/CH2 interface variants, M228T and T247M, increase actin binding, in agreement with our prediction for the interface-localized variants. We also investigated a A119T variant that localizes to an actin-binding surface. This variant was previously reported to decrease actin binding[15]. However, the previously reported decrease in affinity was not statistically significant[15]. Here we clarified the effect of the A119T variant. We showed that the A119T variant, indeed, decreases actin binding, raising the Kd ∼4-fold, with statistical significance. We expect that many of the other 31 variants mapping to direct actin-binding surfaces or the CH1/CH2 interface, will also disrupt actin binding. However, some of the variants are conservative substitutions that may not disrupt structure/function (for example, K38R or R41K). Experimentally confirming the impact of the variants on actin binding is thus important for establishing pathogenicity.

Our thermal denaturation studies revealed that A119T, M228T and T247M are structurally destabilizing. All three variants caused the ABD to begin to unfold at lower temperature relative to wild-type. Unexpectedly, all three variants also caused a loss of cooperative unfolding, with melt curves revealing multiple steps of unfolding for different substructures within the ABD. M228T and T247M showed three unfolding events, and the three different Tm’s were very similar between the two variants. This is consistent with the two variants being localized to the CH1/CH2 interface, and consequently causing similar disruption to ABD structure. In contrast, A119T, buried in CH1, was characterized by two unfolding events, and thus reflects a distinct structural impact on the ABD. Notably, melt curves for all three variants contained an unfolding event with T_m_ of ∼72°C. This shared T_m_ suggests the unfolding of a relatively stable ABD substructure common to all three variants. A loss of cooperative unfolding was not reported previously in thermal denaturation studies for the A119T mutant ABD[15]. Instead the A119T mutant appeared to unfold with a single transition, with a T_m_ of 60.8°C, slightly lower than the 62.3°C reported for wild-type[15]. This difference may reflect greater sensitivity of our CD measurements with a different spectrapolarimeter. The previously characterized A119T ABD also lacked the N-terminal 18 residues; we previously showed that N-terminal residues preceding CH1 impact the stability of the related ABD of β-III-spectrin [27]. Interestingly, loss of cooperative unfolding was not observed for any of ten different missense variants in the related ABD of β-III-spectrin[35]. Most of the β-III-spectrin variants localized to the CH1/CH2 interface, including L253P and T271I, which are at the equivalent residue positions as α-actinin-2 M228T and T247M, respectively. This suggests a fundamental difference in physical coupling of substructures within the β-III-spectrin and α-actinin-2 ABDs.

Our work suggests that altered actin binding underlies pathogenesis for cardiomyopathy mutations in the ABD of α-actinin-2. Prior work in cardiomyocytes showed that A119T has reduced incorporation into Z-disks, is less dynamic at Z-disks and tends to form aggregates outside of Z-disks[15], and were attributed to a loss of actin binding. The T247M variant also cause α-actinin-2 aggregation in iPSC-derived cardiomyocytes[36]. We propose that these aggregates reflect a high-affinity interaction of the T247M mutant with actin filaments outside of the Z-disk structure. Based on the nearly identical impact of the T247M and M228T variants on the structure and function of the ABD, reported here, we expect M228T will show cause similar aggregation and loss of Z-disk incorporation as T247M. Drug-like small molecules have been identified that partially ameliorate high-affinity actin binding of a L253P mutant β-III-spectrin ABD[37]. Drug screening for α-actinin-2 may also identify actin-binding modulators useful in treatment in cardiomyopathy.

## Abbreviations

(ABD): Actin-binding domain
(CD): circular dichroism
(DCM): dilated cardiomyopathy
(HCM): hypertrophic cardiomyopathy
(RCM): restrictive cardiomyopathy

## Data Statement

All data generated and/or analyzed that contributed to this manuscript are available from the corresponding author on reasonable request.

## References

[1] C.W. Tsao, A.W. Aday, Z.I. Almarzooq, C.A.M. Anderson, P. Arora, C.L. Avery, C.M. Baker-Smith, A.Z. Beaton, A.K. Boehme, A.E. Buxton, Y. Commodore-Mensah, M.S.V. Elkind, K.R. Evenson, C. Eze-Nliam, S. Fugar, G. Generoso, D.G. Heard, S. Hiremath, J.E. Ho, R. Kalani, D.S. Kazi, D. Ko, D.A. Levine, J. Liu, J. Ma, J.W. Magnani, E.D. Michos, M.E. Mussolino, S.D. Navaneethan, N.I. Parikh, R. Poudel, M. Rezk-Hanna, G.A. Roth, N.S. Shah, M.P. St-Onge, E.L. Thacker, S.S. Virani, J.H. Voeks, N.Y. Wang, N.D. Wong, S.S. Wong, K. Yaffe, S.S. Martin, E. American Heart Association Council on, C. Prevention Statistics, S. Stroke Statistics, Heart Disease and Stroke Statistics-2023 Update: A Report From the American Heart Association, Circulation, 147 (2023) e93–e621.

[2] R.E. Hershberger, D.J. Hedges, A. Morales, Dilated cardiomyopathy: the complexity of a diverse genetic architecture, Nat Rev Cardiol, 10 (2013) 531–547.

[3] B.J. Maron, J.M. Gardin, J.M. Flack, S.S. Gidding, T.T. Kurosaki, D.E. Bild, Prevalence of hypertrophic cardiomyopathy in a general population of young adults. Echocardiographic analysis of 4111 subjects in the CARDIA Study. Coronary Artery Risk Development in (Young) Adults, Circulation, 92 (1995) 785–789.

[4] E. Muchtar, L.A. Blauwet, M.A. Gertz, Restrictive Cardiomyopathy: Genetics, Pathogenesis, Clinical Manifestations, Diagnosis, and Therapy, Circ Res, 121 (2017) 819–837.

[5] J.A. Towbin, Inherited cardiomyopathies, Circ J, 78 (2014) 2347–2356.

[6] Z.D. Goff, H. Calkins, Sudden death related cardiomyopathies - Hypertrophic cardiomyopathy, Prog Cardiovasc Dis, 62 (2019) 212–216.

[7] B. Mohapatra, S. Jimenez, J.H. Lin, K.R. Bowles, K.J. Coveler, J.G. Marx, M.A. Chrisco, R.T. Murphy, P.R. Lurie, R.J. Schwartz, P.M. Elliott, M. Vatta, W. McKenna, J.A. Towbin, N.E. Bowles, Mutations in the muscle LIM protein and alpha-actinin-2 genes in dilated cardiomyopathy and endocardial fibroelastosis, Mol Genet Metab, 80 (2003) 207–215.

[8] C. Chiu, R.D. Bagnall, J. Ingles, L. Yeates, M. Kennerson, J.A. Donald, M. Jormakka, J.M. Lind, C. Semsarian, Mutations in alpha-actinin-2 cause hypertrophic cardiomyopathy: a genomewide analysis, J Am Coll Cardiol, 55 (2010) 1127–1135.

[9] L.L. Fan, H. Huang, J.Y. Jin, J.J. Li, Y.Q. Chen, R. Xiang, Whole-Exome Sequencing Identifies a Novel Mutation (p.L320R) of Alpha-Actinin 2 in a Chinese Family with Dilated Cardiomyopathy and Ventricular Tachycardia, Cytogenet Genome Res, 157 (2019) 148–152.

[10] M.J. Landrum, J.M. Lee, M. Benson, G.R. Brown, C. Chao, S. Chitipiralla, B. Gu, J. Hart, D. Hoffman, W. Jang, K. Karapetyan, K. Katz, C. Liu, Z. Maddipatla, A. Malheiro, K. McDaniel, M. Ovetsky, G. Riley, G. Zhou, J.B. Holmes, B.L. Kattman, D.R. Maglott, ClinVar: improving access to variant interpretations and supporting evidence, Nucleic Acids Res, 46 (2018) D1062–D1067.

[11] F. Girolami, M. Iascone, B. Tomberli, S. Bardi, M. Benelli, G. Marseglia, C. Pescucci, L. Pezzoli, M.E. Sana, C. Basso, N. Marziliano, P.A. Merlini, A. Fornaro, F. Cecchi, F. Torricelli, I. Olivotto, Novel alpha-actinin 2 variant associated with familial hypertrophic cardiomyopathy and juvenile atrial arrhythmias: a massively parallel sequencing study, Circ Cardiovasc Genet, 7 (2014) 741–750.

[12] A.H. Beggs, T.J. Byers, J.H. Knoll, F.M. Boyce, G.A. Bruns, L.M. Kunkel, Cloning and characterization of two human skeletal muscle alpha-actinin genes located on chromosomes 1 and 11, J Biol Chem, 267 (1992) 9281–9288.

[13] V. Gupta, M. Discenza, J.R. Guyon, L.M. Kunkel, A.H. Beggs, alpha-Actinin-2 deficiency results in sarcomeric defects in zebrafish that cannot be rescued by alpha-actinin-3 revealing functional differences between sarcomeric isoforms, FASEB J, 26 (2012) 1892–1908.

[14] A. Ribeiro Ede, Jr., N. Pinotsis, A. Ghisleni, A. Salmazo, P.V. Konarev, J. Kostan, B. Sjoblom, C. Schreiner, A.A. Polyansky, E.A. Gkougkoulia, M.R. Holt, F.L. Aachmann, B. Zagrovic, E. Bordignon, K.F. Pirker, D.I. Svergun, M. Gautel, K. Djinovic-Carugo, The structure and regulation of human muscle alpha-actinin, Cell, 159 (2014) 1447–1460.

[15] N.J. Haywood, M. Wolny, B. Rogers, C.H. Trinh, Y. Shuping, T.A. Edwards, M. Peckham, Hypertrophic cardiomyopathy mutations in the calponin-homology domain of ACTN2 affect actin binding and cardiomyocyte Z-disc incorporation, Biochem J, 473 (2016) 2485–2493.

[16] M. Prondzynski, M.D. Lemoine, A.T. Zech, A. Horvath, V. Di Mauro, J.T. Koivumaki, N. Kresin, J. Busch, T. Krause, E. Kramer, S. Schlossarek, M. Spohn, F.W. Friedrich, J. Munch, S.D. Laufer, C. Redwood, A.E. Volk, A. Hansen, G. Mearini, D. Catalucci, C. Meyer, T. Christ, M. Patten, T. Eschenhagen, L. Carrier, Disease modeling of a mutation in alpha-actinin 2 guides clinical therapy in hypertrophic cardiomyopathy, EMBO Mol Med, 11 (2019) e11115.

[17] R.K. Liem, Cytoskeletal Integrators: The Spectrin Superfamily, Cold Spring Harb Perspect Biol, 8 (2016).

[18] A.W. Avery, J. Crain, D.D. Thomas, T.S. Hays, A human beta-III-spectrin spinocerebellar ataxia type 5 mutation causes high-affinity F-actin binding, Sci Rep, 6 (2016) 21375.

[19] L.Z. Liu, M. Ren, M. Li, Y.T. Ren, B. Sun, X.S. Sun, S.Y. Chen, S.Y. Li, X.S. Huang, A Novel Missense Mutation in the Spectrin Beta Nonerythrocytic 2 Gene Likely Associated with Spinocerebellar Ataxia Type 5, Chin Med J (Engl), 129 (2016) 2516–2517.

[20] J.M. Kaplan, S.H. Kim, K.N. North, H. Rennke, L.A. Correia, H.Q. Tong, B.J. Mathis, J.C. Rodriguez-Perez, P.G. Allen, A.H. Beggs, M.R. Pollak, Mutations in ACTN4, encoding alphaactinin-4, cause familial focal segmental glomerulosclerosis, Nat Genet, 24 (2000) 251–256.

[21] A. Weins, P. Kenlan, S. Herbert, T.C. Le, I. Villegas, B.S. Kaplan, G.B. Appel, M.R. Pollak, Mutational and Biological Analysis of alpha-actinin-4 in focal segmental glomerulosclerosis, J Am Soc Nephrol, 16 (2005) 3694–3701.

[22] A.C. Murphy, A.J. Lindsay, M.W. McCaffrey, K. Djinovic-Carugo, P.W. Young, Congenital macrothrombocytopenia-linked mutations in the actin-binding domain of alphaactinin-1 enhance F-actin association, FEBS Lett, 590 (2016) 685–695.

[23] A.R. Clark, G.M. Sawyer, S.P. Robertson, A.J. Sutherland-Smith, Skeletal dysplasias due to filamin A mutations result from a gain-of-function mechanism distinct from allelic neurological disorders, Hum Mol Genet, 18 (2009) 4791–4800.

[24] G.M. Sawyer, A.R. Clark, S.P. Robertson, A.J. Sutherland-Smith, Disease-associated substitutions in the filamin B actin binding domain confer enhanced actin binding affinity in the absence of major structural disturbance: Insights from the crystal structures of filamin B actin binding domains, J Mol Biol, 390 (2009) 1030–1047.

[25] R.M. Duff, V. Tay, P. Hackman, G. Ravenscroft, C. McLean, P. Kennedy, A. Steinbach, W. Schoffler, P.F.M. van der Ven, D.O. Furst, J. Song, K. Djinovic-Carugo, S. Penttila, O. Raheem, K. Reardon, A. Malandrini, S. Gambelli, M. Villanova, K.J. Nowak, D.R. Williams, J.E. Landers, R.H. Brown, Jr., B. Udd, N.G. Laing, Mutations in the N-terminal actin-binding domain of filamin C cause a distal myopathy, Am J Hum Genet, 88 (2011) 729–740.

[26] D.M. Henderson, A. Lee, J.M. Ervasti, Disease-causing missense mutations in actin binding domain 1 of dystrophin induce thermodynamic instability and protein aggregation, Proc Natl Acad Sci U S A, 107 (2010) 9632–9637.

[27] A.W. Avery, M.E. Fealey, F. Wang, A. Orlova, A.R. Thompson, D.D. Thomas, T.S. Hays, E.H. Egelman, Structural basis for high-affinity actin binding revealed by a beta-III-spectrin SCA5 missense mutation, Nat Commun, 8 (2017) 1350.

[28] V.E. Galkin, A. Orlova, A. Salmazo, K. Djinovic-Carugo, E.H. Egelman, Opening of tandem calponin homology domains regulates their affinity for F-actin, Nat Struct Mol Biol, 17 (2010) 614–616.

[29] A.T.L. Zech, M. Prondzynski, S.R. Singh, E. Orthey, E. Alizoti, J. Busch, A. Madsen, C.S. Behrens, G. Mearini, M.D. Lemoine, E. Kramer, D. Mosqueira, S. Virdi, D. Indenbirken, M. Depke, M. Gesell Salazar, U. Volker, I. Braren, W.T. Pu, T. Eschenhagen, E. Hammer, S. Schlossarek, L. Carrier, ACTN2 missense variant causes proteopathy in human iPSC-derived cardiomyocytes, bioRxiv, (2021).

[30] S. Broadway-Stringer, H. Jiang, K. Wadmore, C. Hooper, G. Douglas, V. Steeples, A.J. Azad, E. Singer, J.S. Reyat, F. Galatik, E. Ehler, P. Bennett, J.I. Kalisch-Smith, D.B. Sparrow, B. Davies, K. Djinovic-Carugo, M. Gautel, H. Watkins, K. Gehmlich, Insights into the Role of a Cardiomyopathy-Causing Genetic Variant in ACTN2, Cells, 12 (2023).

[31] D.V. Iwamoto, A. Huehn, B. Simon, C. Huet-Calderwood, M. Baldassarre, C.V. Sindelar, D.A. Calderwood, Structural basis of the filamin A actin-binding domain interaction with Factin, Nat Struct Mol Biol, 25 (2018) 918–927.

[32] A. Kumari, S. Kesarwani, M.G. Javoor, K.R. Vinothkumar, M. Sirajuddin, Structural insights into actin filament recognition by commonly used cellular actin markers, EMBO J, 39 (2020) e104006.

[33] S. Legardinier, B. Legrand, C. Raguenes-Nicol, A. Bondon, S. Hardy, C. Tascon, E. Le Rumeur, J.F. Hubert, A Two-amino Acid Mutation Encountered in Duchenne Muscular Dystrophy Decreases Stability of the Rod Domain 23 (R23) Spectrin-like Repeat of Dystrophin, J Biol Chem, 284 (2009) 8822–8832.

[34] S.A. Denha, A.E. Atang, T.S. Hays, A.W. Avery, β-III-spectrin N-terminus is required for high-affinity actin binding and SCA5 neurotoxicity, Sci Rep, In Press (2022).

[35] A.E. Atang, A.R. Keller, S.A. Denha, A.W. Avery, Increased actin binding is a shared molecular consequence of numerous spinocerebellar ataxia mutations in beta-III-spectrin, bioRxiv, (Preprint) (2023).

[36] A.T.L. Zech, M. Prondzynski, S.R. Singh, N. Pietsch, E. Orthey, E. Alizoti, J. Busch, A. Madsen, C.S. Behrens, M. Meyer-Jens, G. Mearini, M.D. Lemoine, E. Kramer, D. Mosqueira, S. Virdi, D. Indenbirken, M. Depke, M.G. Salazar, U. Volker, I. Braren, W.T. Pu, T. Eschenhagen, E. Hammer, S. Schlossarek, L. Carrier, ACTN2 Mutant Causes Proteopathy in Human iPSCDerived Cardiomyocytes, Cells, 11 (2022).

[37] P. Guhathakurta, R.T. Rebbeck, S.A. Denha, A.R. Keller, A.L. Carter, A.E. Atang, B. Svensson, D.D. Thomas, T.S. Hays, A.W. Avery, Early-phase drug discovery of beta-III-spectrin actin-binding modulators for treatment of spinocerebellar ataxia type 5, J Biol Chem, 299 (2023) 102956.

